## Supplementary material for "Modelling spatially autocorrelated detection probabilities in spatial capture-recapture using random effects": Supp. material

**Running headline**: Spatially heterogeneous detectability in SCR

### S1. Tables of posterior estimates and effective sample size (ESS) of different models

**Table S1**: **SCR**: Relative bias (RB, median and 2.5% and 97.5% quantiles), coefficient of variation (CV, median and 2.5% and 97.5% quantiles), and coverage probability of 95% CI of the population size ($N$) and the spatial scale parameter of the half-normal detection function ($\sigma$) across each scenario.

| Scenario | $\eta$ | $\phi$ | RB | | | CV | | | Coverage prob. |
|---|---|---|---|---|---|---|---|---|---|
| | | | Median | 2.5% Quantile | 97.5% Quantile | Median | 2.5% Quantile | 97.5% Quantile | |
| $N$ | | | | | | | | | |
| Continuous | | | | | | | | | |
| 1 | 0.100 | 1 | −0.027 | −0.105 | 0.086 | 0.054 | 0.047 | 0.060 | 0.96 |
| 2 | 0.100 | 0.050 | −0.095 | −0.275 | 0.048 | 0.054 | 0.033 | 0.143 | 0.65 |
| 3 | 0.300 | 1 | 0.005 | −0.060 | 0.070 | 0.033 | 0.031 | 0.034 | 0.94 |
| 4 | 0.300 | 0.050 | −0.025 | −0.116 | 0.044 | 0.033 | 0.027 | 0.046 | 0.79 |
| 5 | 0.600 | 1 | 0.006 | −0.037 | 0.055 | 0.028 | 0.027 | 0.029 | 0.97 |
| 6 | 0.600 | 0.050 | −0.006 | −0.061 | 0.051 | 0.028 | 0.026 | 0.031 | 0.97 |
| Categorical | | | | | | | | | |
| 7 | 0.100 | 1 | −0.032 | −0.185 | 0.271 | 0.137 | 0.111 | 0.166 | 0.99 |
| 8 | 0.100 | 0.050 | −0.295 | −0.438 | −0.040 | 0.113 | 0.092 | 0.138 | 0.20 |
| 9 | 0.300 | 1 | −0.022 | −0.125 | 0.087 | 0.049 | 0.045 | 0.053 | 0.91 |
| 10 | 0.300 | 0.050 | −0.290 | −0.409 | −0.142 | 0.047 | 0.043 | 0.051 | 0 |
| $\sigma$ | | | | | | | | | |
| Continuous | | | | | | | | | |
| 1 | 0.100 | 1 | −0.022 | −0.094 | 0.049 | 0.037 | 0.033 | 0.042 | 0.90 |
| 2 | 0.100 | 0.050 | −0.012 | −0.098 | 0.068 | 0.037 | 0.019 | 0.086 | 0.94 |
| 3 | 0.300 | 1 | −0.007 | −0.042 | 0.032 | 0.019 | 0.017 | 0.020 | 0.92 |
| 4 | 0.300 | 0.050 | −0.004 | −0.046 | 0.030 | 0.019 | 0.012 | 0.031 | 0.93 |
| 5 | 0.600 | 1 | −0.001 | −0.031 | 0.026 | 0.013 | 0.012 | 0.014 | 0.90 |
| 6 | 0.600 | 0.050 | −0.001 | −0.024 | 0.023 | 0.013 | 0.010 | 0.016 | 0.95 |
| Categorical | | | | | | | | | |
| 7 | 0.100 | 1 | −0.026 | −0.195 | 0.170 | 0.085 | 0.073 | 0.103 | 0.88 |
| 8 | 0.100 | 0.050 | −0.032 | −0.185 | 0.144 | 0.073 | 0.062 | 0.085 | 0.82 |
| 9 | 0.300 | 1 | −0.020 | −0.103 | 0.041 | 0.034 | 0.031 | 0.037 | 0.81 |
| 10 | 0.300 | 0.050 | −0.038 | −0.100 | 0.022 | 0.031 | 0.028 | 0.034 | 0.72 |

**Table S2**: **RE** ($4 \times 4$): Relative bias (RB, median and 2.5% and 97.5% quantiles), coefficient of variation (CV, median and 2.5% and 97.5% quantiles), and coverage probability of 95% CI of the population size ($N$) and the spatial scale parameter of the half-normal detection function ($\sigma$) across each scenario. Here '$4 \times 4$' refers to the level to aggregation in the random effects (Section 3.2, main text).

| Scenario | $\eta$ | $\phi$ | RB | | | CV | | | Coverage prob. |
|---|---|---|---|---|---|---|---|---|---|
| | | | Median | 2.5% Quantile | 97.5% Quantile | Median | 2.5% Quantile | 97.5% Quantile | |
| $N$ | | | | | | | | | |
| Continuous | | | | | | | | | |
| 1 | 0.100 | 1 | 0.032 | −0.056 | 0.133 | 0.058 | 0.051 | 0.065 | 0.96 |
| 2 | 0.100 | 0.050 | −0.017 | −0.141 | 0.152 | 0.062 | 0.035 | 0.149 | 0.94 |
| 3 | 0.300 | 1 | 0.015 | −0.048 | 0.077 | 0.034 | 0.032 | 0.036 | 0.92 |
| 4 | 0.300 | 0.050 | −0.003 | −0.082 | 0.065 | 0.036 | 0.028 | 0.052 | 0.94 |
| 5 | 0.600 | 1 | 0.010 | −0.035 | 0.065 | 0.029 | 0.028 | 0.030 | 0.95 |
| 6 | 0.600 | 0.050 | 0.002 | −0.049 | 0.053 | 0.029 | 0.026 | 0.033 | 0.99 |
| Categorical | | | | | | | | | |
| 7 | 0.100 | 1 | 0.047 | −0.118 | 0.349 | 0.139 | 0.115 | 0.162 | 0.94 |
| 8 | 0.100 | 0.050 | −0.065 | −0.296 | 0.168 | 0.133 | 0.117 | 0.153 | 0.90 |
| 9 | 0.300 | 1 | 0.037 | −0.082 | 0.131 | 0.054 | 0.049 | 0.057 | 0.94 |
| 10 | 0.300 | 0.050 | −0.166 | −0.291 | −0.016 | 0.063 | 0.055 | 0.071 | 0.29 |
| $\sigma$ | | | | | | | | | |
| Continuous | | | | | | | | | |
| 1 | 0.100 | 1 | −0.010 | −0.094 | 0.059 | 0.038 | 0.034 | 0.043 | 0.93 |
| 2 | 0.100 | 0.050 | −0.008 | −0.083 | 0.080 | 0.038 | 0.019 | 0.090 | 0.94 |
| 3 | 0.300 | 1 | −0.008 | −0.040 | 0.030 | 0.019 | 0.018 | 0.021 | 0.93 |
| 4 | 0.300 | 0.050 | −0.003 | −0.040 | 0.035 | 0.019 | 0.012 | 0.033 | 0.92 |
| 5 | 0.600 | 1 | −0.007 | −0.039 | 0.022 | 0.013 | 0.012 | 0.014 | 0.86 |
| 6 | 0.600 | 0.050 | −0.005 | −0.028 | 0.019 | 0.013 | 0.009 | 0.016 | 0.94 |
| Categorical | | | | | | | | | |
| 7 | 0.100 | 1 | −0.012 | −0.196 | 0.212 | 0.087 | 0.076 | 0.104 | 0.89 |
| 8 | 0.100 | 0.050 | −0.007 | −0.160 | 0.186 | 0.077 | 0.066 | 0.091 | 0.86 |
| 9 | 0.300 | 1 | −0.008 | −0.092 | 0.047 | 0.035 | 0.031 | 0.038 | 0.89 |
| 10 | 0.300 | 0.050 | −0.015 | −0.075 | 0.044 | 0.032 | 0.029 | 0.036 | 0.92 |

**Table S3**: **SARE** ($4 \times 4$): Relative bias (RB, median and 2.5% and 97.5% quantiles), coefficient of variation (CV, median and 2.5% and 97.5% quantiles), and coverage probability of 95% CI of the population size ($N$) and the spatial scale parameter of the half-normal detection function ($\sigma$) across each scenario. Here '$4 \times 4$' refers to the level to aggregation in the random effects (Section 3.2, main text).

| Scenario | $\eta$ | $\phi$ | RB | | | CV | | | Coverage prob. |
|---|---|---|---|---|---|---|---|---|---|
| | | | Median | 2.5% Quantile | 97.5% Quantile | Median | 2.5% Quantile | 97.5% Quantile | |
| $N$ | | | | | | | | | |
| Continuous | | | | | | | | | |
| 1 | 0.100 | 1 | −0.014 | −0.077 | 0.096 | 0.055 | 0.049 | 0.064 | 1 |
| 2 | 0.100 | 0.050 | −0.004 | −0.130 | 0.189 | 0.055 | 0.035 | 0.138 | 0.94 |
| 3 | 0.300 | 1 | 0.020 | −0.061 | 0.079 | 0.034 | 0.031 | 0.035 | 0.90 |
| 4 | 0.300 | 0.050 | 0.001 | −0.082 | 0.071 | 0.036 | 0.028 | 0.051 | 0.93 |
| 5 | 0.600 | 1 | 0.016 | −0.024 | 0.051 | 0.028 | 0.028 | 0.029 | 0.97 |
| 6 | 0.600 | 0.050 | 0.0002 | −0.051 | 0.049 | 0.029 | 0.026 | 0.033 | 0.99 |
| Categorical | | | | | | | | | |
| 7 | 0.100 | 1 | 0.064 | −0.017 | 0.242 | 0.132 | 0.123 | 0.142 | 1 |
| 8 | 0.100 | 0.050 | 0.002 | −0.235 | 0.229 | 0.131 | 0.117 | 0.146 | 0.91 |
| 9 | 0.300 | 1 | 0.032 | −0.074 | 0.101 | 0.053 | 0.048 | 0.057 | 0.97 |
| 10 | 0.300 | 0.050 | −0.091 | −0.219 | 0.035 | 0.066 | 0.057 | 0.074 | 0.77 |
| $\sigma$ | | | | | | | | | |
| Continuous | | | | | | | | | |
| 1 | 0.100 | 1 | −0.006 | −0.080 | 0.049 | 0.035 | 0.030 | 0.039 | 0.89 |
| 2 | 0.100 | 0.050 | 0.001 | −0.077 | 0.074 | 0.031 | 0.018 | 0.078 | 0.94 |
| 3 | 0.300 | 1 | 0.003 | −0.037 | 0.034 | 0.019 | 0.017 | 0.020 | 0.95 |
| 4 | 0.300 | 0.050 | 0.001 | −0.038 | 0.041 | 0.019 | 0.012 | 0.031 | 0.94 |
| 5 | 0.600 | 1 | −0.001 | −0.032 | 0.020 | 0.013 | 0.012 | 0.013 | 0.90 |
| 6 | 0.600 | 0.050 | 0.001 | −0.027 | 0.022 | 0.013 | 0.010 | 0.016 | 0.94 |
| Categorical | | | | | | | | | |
| 7 | 0.100 | 1 | −0.024 | −0.204 | 0.052 | 0.076 | 0.067 | 0.088 | 0.60 |
| 8 | 0.100 | 0.050 | −0.013 | −0.160 | 0.189 | 0.074 | 0.062 | 0.089 | 0.85 |
| 9 | 0.300 | 1 | −0.010 | −0.104 | 0.052 | 0.033 | 0.031 | 0.038 | 0.87 |
| 10 | 0.300 | 0.050 | −0.011 | −0.075 | 0.046 | 0.032 | 0.028 | 0.036 | 0.92 |

**Table S4**: **FM** ($4\times4$): Relative bias (RB, median and 2.5% and 97.5% quantiles), coefficient of variation (CV, median and 2.5% and 97.5% quantiles), and coverage probability of 95% CI of the population size ($N$) and the spatial scale parameter of the half-normal detection function ($\sigma$) across each scenario. Here '$4 \times 4$' refers to the level to aggregation in the random effects (Section 3.2, main text).

| Scenario | $\eta$ | $\phi$ | RB | | | CV | | | Coverage prob. |
|---|---|---|---|---|---|---|---|---|---|
| | | | Median | 2.5% Quantile | 97.5% Quantile | Median | 2.5% Quantile | 97.5% Quantile | |
| $N$ | | | | | | | | | |
| Continuous | | | | | | | | | |
| 1 | 0.100 | 1 | −0.009 | −0.083 | 0.084 | 0.056 | 0.050 | 0.064 | 0.98 |
| 2 | 0.100 | 0.050 | −0.057 | −0.202 | 0.067 | 0.056 | 0.034 | 0.150 | 0.82 |
| 3 | 0.300 | 1 | 0.008 | −0.051 | 0.071 | 0.033 | 0.031 | 0.035 | 0.95 |
| 4 | 0.300 | 0.050 | −0.011 | −0.092 | 0.060 | 0.034 | 0.028 | 0.051 | 0.93 |
| 5 | 0.600 | 1 | 0.008 | −0.037 | 0.057 | 0.028 | 0.027 | 0.029 | 0.96 |
| 6 | 0.600 | 0.050 | −0.0001 | −0.053 | 0.050 | 0.028 | 0.026 | 0.032 | 0.99 |
| Categorical | | | | | | | | | |
| 7 | 0.100 | 1 | −0.021 | −0.184 | 0.299 | 0.137 | 0.115 | 0.169 | 0.97 |
| 8 | 0.100 | 0.050 | −0.112 | −0.352 | 0.125 | 0.137 | 0.120 | 0.161 | 0.84 |
| 9 | 0.300 | 1 | 0.004 | −0.109 | 0.076 | 0.051 | 0.047 | 0.057 | 0.95 |
| 10 | 0.300 | 0.050 | −0.115 | −0.248 | 0.020 | 0.064 | 0.052 | 0.078 | 0.51 |
| $\sigma$ | | | | | | | | | |
| Continuous | | | | | | | | | |
| 1 | 0.100 | 1 | −0.013 | −0.093 | 0.052 | 0.037 | 0.033 | 0.042 | 0.929 |
| 2 | 0.100 | 0.050 | −0.005 | −0.096 | 0.059 | 0.034 | 0.019 | 0.086 | 0.942 |
| 3 | 0.300 | 1 | −0.005 | −0.040 | 0.035 | 0.019 | 0.017 | 0.021 | 0.928 |
| 4 | 0.300 | 0.050 | −0.001 | −0.038 | 0.036 | 0.019 | 0.012 | 0.032 | 0.939 |
| 5 | 0.600 | 1 | −0.001 | −0.030 | 0.025 | 0.013 | 0.012 | 0.014 | 0.896 |
| 6 | 0.600 | 0.050 | 0.001 | −0.024 | 0.022 | 0.013 | 0.010 | 0.016 | 0.960 |
| Categorical | | | | | | | | | |
| 7 | 0.100 | 1 | −0.044 | −0.196 | 0.162 | 0.085 | 0.073 | 0.101 | 0.864 |
| 8 | 0.100 | 0.050 | −0.024 | −0.172 | 0.185 | 0.075 | 0.063 | 0.088 | 0.837 |
| 9 | 0.300 | 1 | −0.017 | −0.096 | 0.042 | 0.034 | 0.031 | 0.037 | 0.850 |
| 10 | 0.300 | 0.050 | −0.018 | −0.079 | 0.039 | 0.032 | 0.028 | 0.036 | 0.909 |

**Table S5**: **FE**: Relative bias (RB, median and 2.5% and 97.5% quantiles), coefficient of variation (CV, median and 2.5% and 97.5% quantiles), and coverage probability of 95% CI of the population size ($N$) and the spatial scale parameter of the half-normal detection function ($\sigma$) across each scenario.

| Scenario | $\eta$ | $\phi$ | RB | | | CV | | | Coverage prob. |
|---|---|---|---|---|---|---|---|---|---|
| | | | Median | 2.5% Quantile | 97.5% Quantile | Median | 2.5% Quantile | 97.5% Quantile | |
| $N$ | | | | | | | | | |
| Continuous | | | | | | | | | |
| 1 | 0.100 | 1 | 0.017 | −0.070 | 0.101 | 0.054 | 0.049 | 0.059 | 0.980 |
| 2 | 0.100 | 0.050 | 0.007 | −0.103 | 0.152 | 0.057 | 0.035 | 0.123 | 0.950 |
| 3 | 0.300 | 1 | 0.010 | −0.055 | 0.070 | 0.033 | 0.031 | 0.035 | 0.960 |
| 4 | 0.300 | 0.050 | 0.008 | −0.067 | 0.083 | 0.035 | 0.028 | 0.050 | 0.920 |
| 5 | 0.600 | 1 | 0.007 | −0.037 | 0.060 | 0.028 | 0.027 | 0.029 | 0.960 |
| 6 | 0.600 | 0.050 | 0.007 | −0.047 | 0.047 | 0.028 | 0.026 | 0.033 | 0.980 |
| Categorical | | | | | | | | | |
| 7 | 0.100 | 1 | 0.033 | −0.150 | 0.292 | 0.121 | 0.104 | 0.138 | 0.950 |
| 8 | 0.100 | 0.050 | 0.057 | −0.129 | 0.323 | 0.113 | 0.100 | 0.127 | 0.950 |
| 9 | 0.300 | 1 | 0.020 | −0.079 | 0.117 | 0.050 | 0.047 | 0.054 | 0.940 |
| 10 | 0.300 | 0.050 | 0.015 | −0.085 | 0.128 | 0.062 | 0.055 | 0.067 | 0.930 |
| $\sigma$ | | | | | | | | | |
| Continuous | | | | | | | | | |
| 1 | 0.100 | 1 | 0.0003 | −0.067 | 0.045 | 0.026 | 0.023 | 0.030 | 0.940 |
| 2 | 0.100 | 0.050 | 0.001 | −0.073 | 0.070 | 0.026 | 0.014 | 0.063 | 0.960 |
| 3 | 0.300 | 1 | 0.002 | −0.021 | 0.027 | 0.014 | 0.012 | 0.015 | 0.960 |
| 4 | 0.300 | 0.050 | 0.001 | −0.029 | 0.030 | 0.014 | 0.009 | 0.023 | 0.960 |
| 5 | 0.600 | 1 | 0.003 | −0.021 | 0.021 | 0.010 | 0.009 | 0.010 | 0.910 |
| 6 | 0.600 | 0.050 | −0.0005 | −0.016 | 0.018 | 0.010 | 0.008 | 0.012 | 0.970 |
| Categorical | | | | | | | | | |
| 7 | 0.100 | 1 | −0.008 | −0.102 | 0.120 | 0.061 | 0.053 | 0.069 | 0.940 |
| 8 | 0.100 | 0.050 | 0.002 | −0.090 | 0.122 | 0.055 | 0.048 | 0.063 | 0.950 |
| 9 | 0.300 | 1 | 0.003 | −0.051 | 0.045 | 0.024 | 0.022 | 0.026 | 0.950 |
| 10 | 0.300 | 0.050 | 0.006 | −0.037 | 0.042 | 0.023 | 0.021 | 0.027 | 0.990 |

**Table S6**: **SCR** and **FE**: Effective sample size (ESS) after burn-in, MCMC efficiency (=ESS/MCMC runtime) and full MCMC run time of a single chain (in minutes). For comparison purposes, we report mean ESS and mean MCMC efficiency (=ESS/MCMC runtime) averaged over each top-level parameters in the model and over each of the converged replicates. For both SCR and FE models, we ran three MCMC chains of 30,000 iterations (burn-in period 12,000).

| Scenario | $\eta$ | $\phi$ | SCR | | | FE | | |
|---|---|---|---|---|---|---|---|---|
| | | | ESS | MCMC efficiency | MCMC run time | ESS | MCMC efficiency | MCMC run time |
| Continuous | | | | | | | | |
| 1 | 0.100 | 1 | 2277 | 1.224 | 17.260 | 10032 | 1.067 | 65.210 |
| 2 | 0.100 | 0.050 | 2369 | 1.312 | 16.520 | 9909 | 1.065 | 65.070 |
| 3 | 0.300 | 1 | 3608 | 1.853 | 18.050 | 12562 | 1.323 | 65.770 |
| 4 | 0.300 | 0.050 | 3691 | 1.922 | 17.790 | 12333 | 1.311 | 65.620 |
| 5 | 0.600 | 1 | 4133 | 2.096 | 18.280 | 13398 | 1.413 | 65.820 |
| 6 | 0.600 | 0.050 | 4215 | 2.145 | 18.210 | 13354 | 1.418 | 65.680 |
| Categorical | | | | | | | | |
| 7 | 0.100 | 1 | 698 | 0.412 | 15.870 | 6160 | 0.876 | 48.910 |
| 8 | 0.100 | 0.050 | 1274 | 0.839 | 14.220 | 6362 | 0.901 | 49.140 |
| 9 | 0.300 | 1 | 2521 | 1.392 | 16.830 | 8453 | 1.176 | 50 |
| 10 | 0.300 | 0.050 | 3697 | 2.312 | 14.890 | 8187 | 1.153 | 49.410 |

**Table S7**: **RE**: Effective sample size (ESS) after burn-in, MCMC efficiency (=ESS/MCMC runtime) and full MCMC run time of a single chain (in minutes). For comparison purposes, we report mean ESS and mean MCMC efficiency (=ESS/MCMC runtime) averaged over each top-level parameters in the model and over each of the converged replicates. We ran three MCMC chains of 100,000 iterations (burn-in period 20,000, with or without aggregation). Scenarios without any converged replicates are indicated by '-'. Here '$1 \times 1$' indicates that the model is fitted without aggregation and '$4 \times 4$' refers to the level to aggregation in the random effects (Section 3.2, main text).

| Scenario | $\eta$ | $\phi$ | RE ($1 \times 1$) | | | RE ($4 \times 4$) | | |
|---|---|---|---|---|---|---|---|---|
| | | | ESS | MCMC efficiency | MCMC run time | ESS | MCMC efficiency | MCMC run time |
| Continuous | | | | | | | | |
| 1 | 0.100 | 1 | 8976 | 0.557 | 112.140 | 5872 | 0.543 | 75.380 |
| 2 | 0.100 | 0.050 | 7525 | 0.486 | 108.690 | 5509 | 0.520 | 73.340 |
| 3 | 0.300 | 1 | 13420 | 0.813 | 115.280 | 9581 | 0.867 | 76.830 |
| 4 | 0.300 | 0.050 | 13671 | 0.847 | 112.710 | 9112 | 0.832 | 76.050 |
| 5 | 0.600 | 1 | 14221 | 0.855 | 115.110 | 11152 | 1.004 | 77.260 |
| 6 | 0.600 | 0.050 | 14775 | 0.901 | 114.360 | 11133 | 1.004 | 77.060 |
| Categorical | | | | | | | | |
| 7 | 0.100 | 1 | 2897 | 0.185 | 109.560 | 2338 | 0.229 | 71.930 |
| 8 | 0.100 | 0.050 | 5166 | 0.360 | 100.490 | 2935 | 0.311 | 66.490 |
| 9 | 0.300 | 1 | 9875 | 0.610 | 112.260 | 6292 | 0.591 | 74.320 |
| 10 | 0.300 | 0.050 | 14092 | 0.961 | 102.290 | 4996 | 0.524 | 66.280 |

**Table S8**: **SARE**: Effective sample size (ESS) after burn-in, MCMC efficiency (=ESS/MCMC runtime) and full MCMC run time of a single chain (in minutes). For comparison purposes, we report mean ESS and mean MCMC efficiency (=ESS/MCMC runtime) averaged over each top-level parameters in the model and over each of the converged replicates. We ran three MCMC chains of 100,000 iterations (burn-in period 20,000, with or without aggregation). Scenarios without any converged replicates are indicated by '-'. Here '$1 \times 1$' indicates that the model is fitted without aggregation and '$4 \times 4$' refers to the level to aggregation in the random effects (Section 3.2, main text).

| Scenario | $\eta$ | $\phi$ | SARE ($1 \times 1$) | | | SARE ($4 \times 4$) | | |
|---|---|---|---|---|---|---|---|---|
| | | | ESS | MCMC efficiency | MCMC run time | ESS | MCMC efficiency | MCMC run time |
| Continuous | | | | | | | | |
| 1 | 0.100 | 1 | - | - | 224.950 | 8803 | 1.028 | 60 |
| 2 | 0.100 | 0.050 | - | - | 221.370 | 6967 | 0.809 | 58.870 |
| 3 | 0.300 | 1 | - | - | 228.440 | 12234 | 1.361 | 62 |
| 4 | 0.300 | 0.050 | - | - | 229.610 | 10924 | 1.262 | 60.180 |
| 5 | 0.600 | 1 | - | - | 230.220 | 13749 | 1.564 | 62.060 |
| 6 | 0.600 | 0.050 | - | - | 231.900 | 12342 | 1.406 | 61.200 |
| Categorical | | | | | | | | |
| 7 | 0.100 | 1 | - | - | 217.490 | 6634 | 0.774 | 57.070 |
| 8 | 0.100 | 0.050 | - | - | 214.990 | 2214 | 0.274 | 56.070 |
| 9 | 0.300 | 1 | - | - | 226.890 | 6108 | 0.699 | 60.540 |
| 10 | 0.300 | 0.050 | - | - | 221.840 | 4495 | 0.544 | 57.470 |

**Table S9**: **FM**: Effective sample size (ESS) after burn-in, MCMC efficiency (=ESS/MCMC runtime) and full MCMC run time of a single chain (in minutes). For comparison purposes, we report mean ESS and mean MCMC efficiency (=ESS/MCMC runtime) averaged over each top-level parameters in the model and over each of the converged replicates. We ran the MCMC for a. 60,000 iterations (burn-in period 12,000) when fitted without aggregation, b. 20,000 iterations (burn-in period 4,000) when fitted with aggregating random effects. Scenarios without any converged replicates are indicated by '-'. Here '$1 \times 1$' and '$4 \times 4$' refer to the level to aggregation in the random effects - '$1 \times 1$' indicating that the model is fitted without aggregation and '$4 \times 4$' indicating that we aggregated the random effects at $4 \times 4$ scale (Section 3.2, main text).

| Scenario | $\eta$ | $\phi$ | FM ($1 \times 1$) | | | FM ($4 \times 4$) | | |
|---|---|---|---|---|---|---|---|---|
| | | | ESS | MCMC efficiency | MCMC run time | ESS | MCMC efficiency | MCMC run time |
| Continuous | | | | | | | | |
| 1 | 0.100 | 1 | 2865 | 0.072 | 283.140 | 1104 | 0.037 | 207.470 |
| 2 | 0.100 | 0.050 | 3258 | 0.086 | 260.920 | 1387 | 0.049 | 197.580 |
| 3 | 0.300 | 1 | 4111 | 0.095 | 292.790 | 1758 | 0.058 | 212.690 |
| 4 | 0.300 | 0.050 | 4312 | 0.106 | 281.680 | 2189 | 0.073 | 208.760 |
| 5 | 0.600 | 1 | 4822 | 0.114 | 294.450 | 1929 | 0.062 | 214.630 |
| 6 | 0.600 | 0.050 | 4820 | 0.119 | 289.410 | 2294 | 0.075 | 212.550 |
| Categorical | | | | | | | | |
| 7 | 0.100 | 1 | - | - | 283.850 | 432 | 0.015 | 210.310 |
| 8 | 0.100 | 0.050 | 1561 | 0.059 | 223.690 | 967 | 0.036 | 188.820 |
| 9 | 0.300 | 1 | 3106 | 0.076 | 288.380 | 1269 | 0.042 | 210.900 |
| 10 | 0.300 | 0.050 | 4386 | 0.134 | 224.860 | 2585 | 0.095 | 189.170 |

### S2. Additional figures

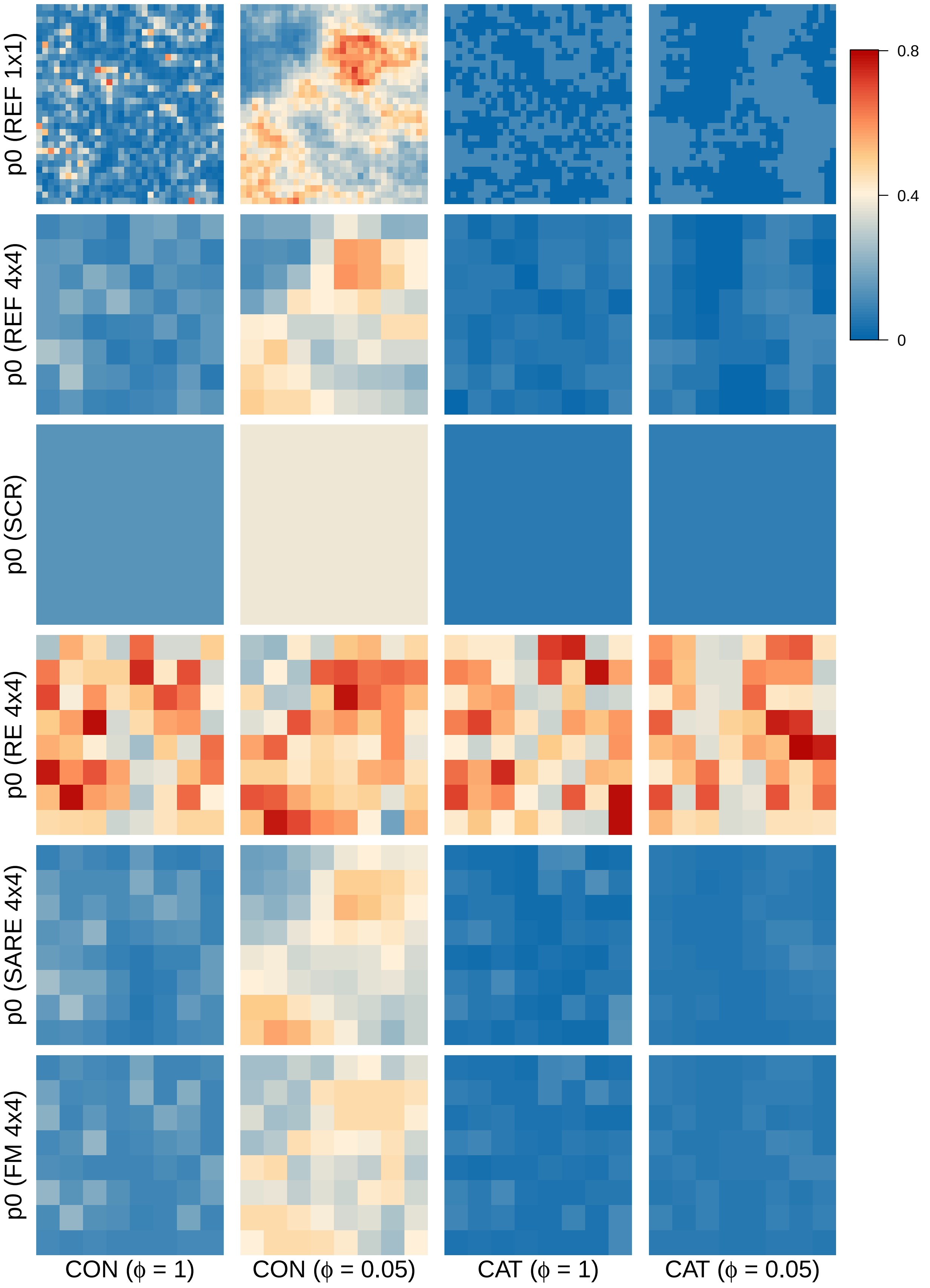


**Figure S1**: Illustration of baseline detection probability surfaces for simulation scenarios under average baseline detection probability $\eta = 0.1$. In rows: simulated baseline detection probability surface ('REF $1 \times 1$'), baseline detection probability surface after averaging the simulated values for each cluster of detectors at $4 \times 4$ scale ('REF $4 \times 4$'), predicted baseline detection probability surface from four models: SCR, RE (aggregation $4 \times 4$), SARE (aggregation $4 \times 4$), FM (aggregation $4 \times 4$). Labels on the $x$-axis refer to scenarios with continuous ("CON") and categorical ("CAT") detector-specific variation in detection probability. Second and fourth columns represent high autocorrelation among detectors (i.e., under $\phi = 0.05$), whereas first and the third column represent intermediate autocorrelation (i.e., under $\phi = 1$). Colors correspond to different values of baseline detection probability.

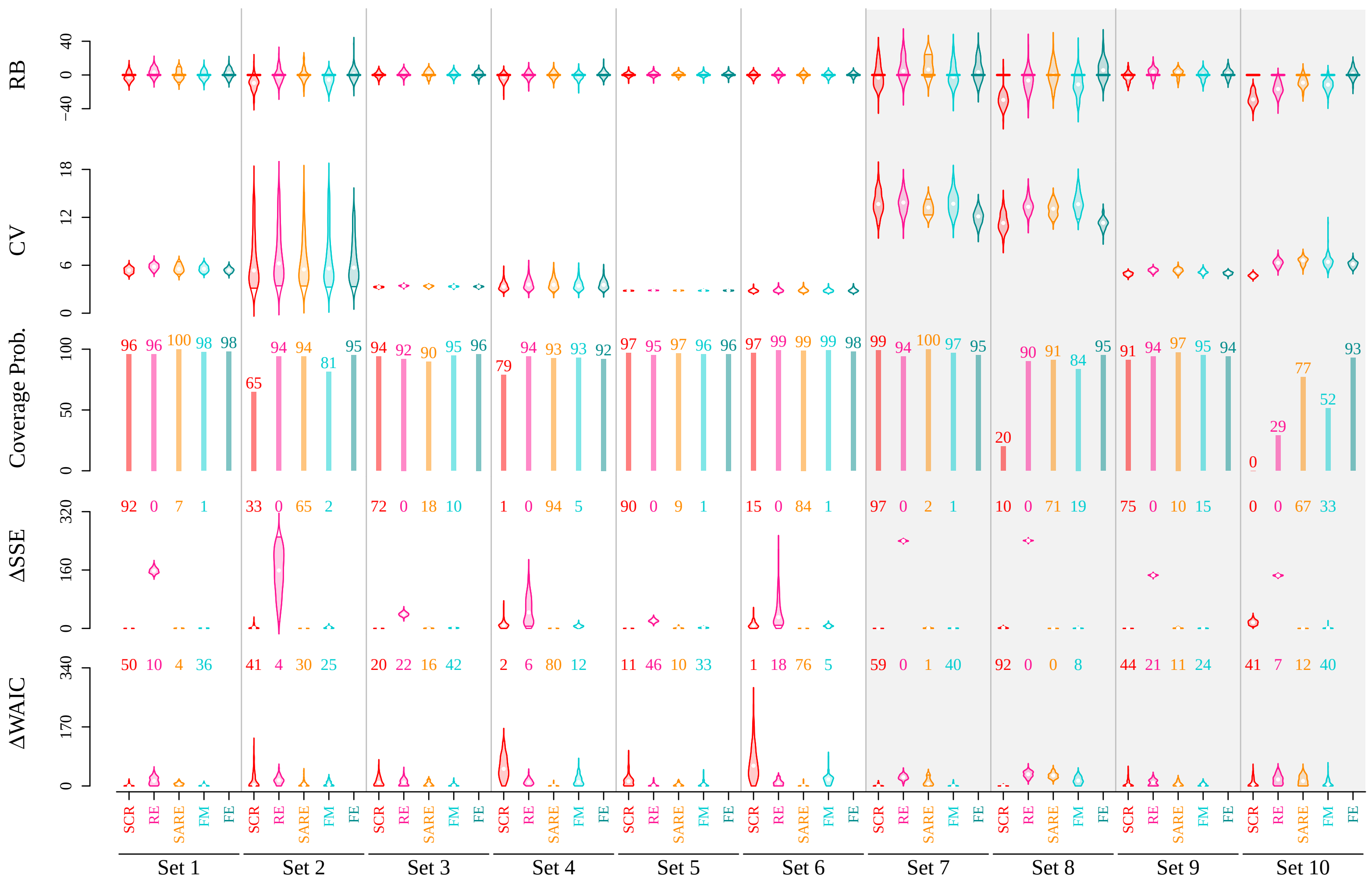


**Figure S2**: Posterior summaries of population size $N$ derived using five models: (i) BASIC-SCR, (ii) SARE (aggregation $4 \times 4$), (iii) RE (aggregation $4 \times 4$), (iv) FM (aggregation $4 \times 4$), (v) FE. Results compare relative bias (RB, in %), coefficient of variation (CV, in %), and coverage probability (in %) of 95% CI for different sets of simulation scenarios. Violins represent the distribution of RB/CV from 100 simulations. Set numbers on the $x$-axis refer to the serial number of the simulation scenario (as shown in the tables). Grey shaded background represents scenarios with categorical detector specific variation, whereas white background represents continuous variation. The figures were based on models that met convergence criteria (Section 3.3, main text).

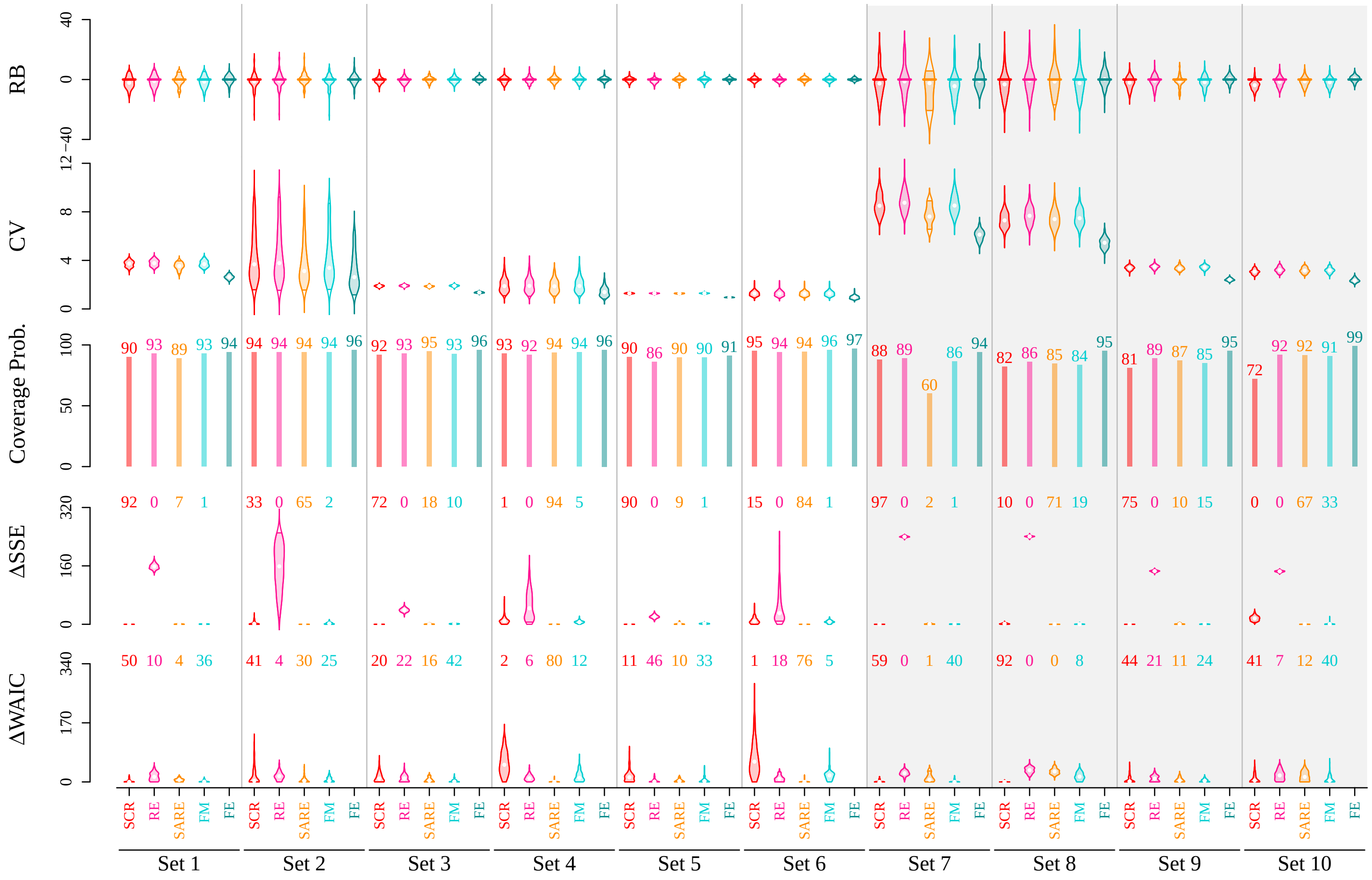


**Figure S3**: Posterior summaries of $\sigma$ derived using five models: (i) BASIC-SCR, (ii) SARE (aggregation $4 \times 4$), (iii) RE (aggregation $4 \times 4$), (iv) FM (aggregation $4 \times 4$), (v) FE. Results compare relative bias (RB, in %), coefficient of variation (CV, in %), and coverage probability (in %) of 95% CI for different sets of simulation scenarios. Violins represent the distribution of RB/CV from 100 simulations. Set numbers on the $x$-axis refer to the serial number of the simulation scenario (as shown in the tables). Grey shaded background represents scenarios with categorical detector specific variation, whereas white background represents continuous variation. The figures were based on models that met convergence criteria (Section 3.3, main text).

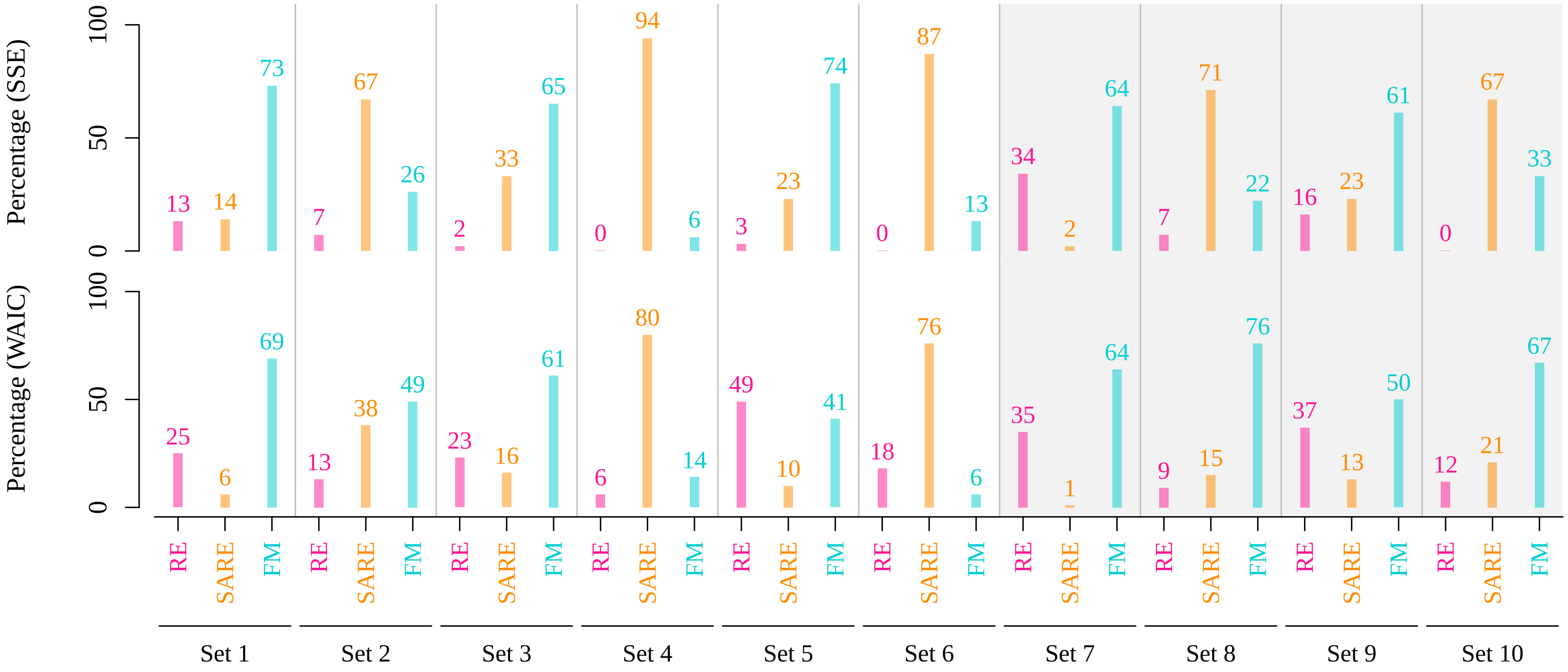


**Figure S4**: Barplot showing percentage of replicates where SSE of predicted baseline detection probability surface and WAIC from a fitted model is lowest among its competitors. Here we considered only the three SCR-GLMMs, namely RE, SARE and FM. Set numbers on the $x$-axis refer to the serial number of the simulation scenario (as shown in the tables). Grey shaded background represents scenarios with categorical detector specific variation, whereas white background represents continuous variation. The figures were based on models that met convergence criteria and the MCMC chains were of different lengths for different models (Section 3.3, main text).
